# Nociceptive neurons interact directly with gastric cancer cells via a CGRP/Ramp1 axis to promote tumor progression

**DOI:** 10.1101/2024.03.04.583209

**Authors:** Xiaofei Zhi, Feijing Wu, Jin Qian, Yosuke Ochiai, Guodong Lian, Ermanno Malagola, Duan Chen, Sandra W Ryeom, Timothy C Wang

## Abstract

Cancer cells have been shown to exploit neurons to modulate their survival and growth, including through establishment of neural circuits within the central nervous system (CNS) ^1–3^. Here, we report a distinct pattern of cancer-nerve interactions between the peripheral nervous system (PNS) and gastric cancer (GC). In multiple GC mouse models, nociceptive nerves demonstrated the greatest degree of nerve expansion in an NGF-dependent manner. Neural tracing identified CGRP+ peptidergic neurons as the primary gastric sensory neurons. Three-dimensional co-culture models showed that sensory neurons directly connect with gastric cancer spheroids through synapse-like structures. Chemogenetic activation of sensory neurons induced the release of calcium into the cytoplasm of cancer cells, promoting tumor growth and metastasis. Pharmacological ablation of sensory neurons or treatment with CGRP inhibitors suppressed tumor growth and extended survival. Depolarization of gastric tumor membranes through *in vivo* optogenetic activation led to enhanced calcium flux in nodose ganglia and CGRP release, defining a cancer cell-peptidergic neuronal circuit. Together, these findings establish the functional connectivity between cancer and sensory neurons, identifying this pathway as a potential therapeutic target.

## Introduction

The peripheral nervous system (PNS) is a critical part of the tumor microenvironment, contributing substantially to tumorigenesis and tumor progression ^4^. In central nervous system (CNS) tumours, cancer cells interact directly with surrounding neurons through electrical and chemical communications ^1–3,5,6^. The interactions in extracranial tumors are less clear. While the full range of neurotransmitters are present in CNS, including amino acid neurotransmitters, the PNS comprises both autonomic and sensory nerves, and thus relies heavily on acetylcholine, norepinephrine, and a variety of neuropeptides as neurotransmitters ^7^. Pain-initiating sensory neurons have been shown to play indirect roles in solid tumor growth through suppression of cancer immunosurveillance ^8^ and induction of cytoprotective autophagy ^9^. Brain tumors such as gliomas originate from neural/glial cells that express pre- and post-synaptic genes and thus display true synapses ^2,3^. In contrast, analysis of extra-cranial solid tumors to date have provided no evidence to support the presence of synapse-like structure.

Interestingly, there appears to be organ-specific variation in the type of tumor innervation ^10^. The tropism of peripheral nerve recruitment highlights the importance of studying tumor innervation in an organ-specific manner. While we previously found that vagal denervation markedly reduced the growth of gastric cancer ^11^, sensory nerves are a major component of vagal axons ^12^, raising questions regarding the role of sensory nerves in gastric carcinogenesis. Here, we report the innervation profile of normal stomach and gastric cancer (GC) and reveal a direct interaction between gastric cancer cells and peripheral nociceptive neurons.

## Results

### Sensory nerves show the greatest degree of nerve expansion within gastric cancer in an NGF-dependent manner

We studied the neural innervation of GC using four different GC mouse models (MNU model ^13^, *Cck2r*-CreERT; *Kras*^G12D^ model ^14^, *Atp4b*-Cre; *Cdh1*^fl/fl^; *Kras*^G12D^; *Trp53* ^fl/fl^; YFP model ^15^, and a syngeneic orthotopic allograft model), initially through immunostaining. While VAChT+ parasympathetic nerves were the most abundant nerves innervating the normal stomach, CGRP+ sensory nerves showed the greatest degree of expansion throughout the glands in all four GC mouse models. (**Fig. 1a-b and Extended Data Fig. 1a-b**). Gastric orthotopic tumors were primarily innervated by CGRP+ nociceptive nerves, while subcutaneous tumors showed a greater abundance of NF200+ fibers (**Extended Data Fig. 1c**), indicating site-and/or organ-specific variation in innervation. Studies of human GC tumors using tissue microarrays confirmed that both intestinal-and diffuse-type gastric cancer showed a higher density of CGRP+ nerves compared to normal human stomach (**Fig. 1c-d**).

**Figure 1.**
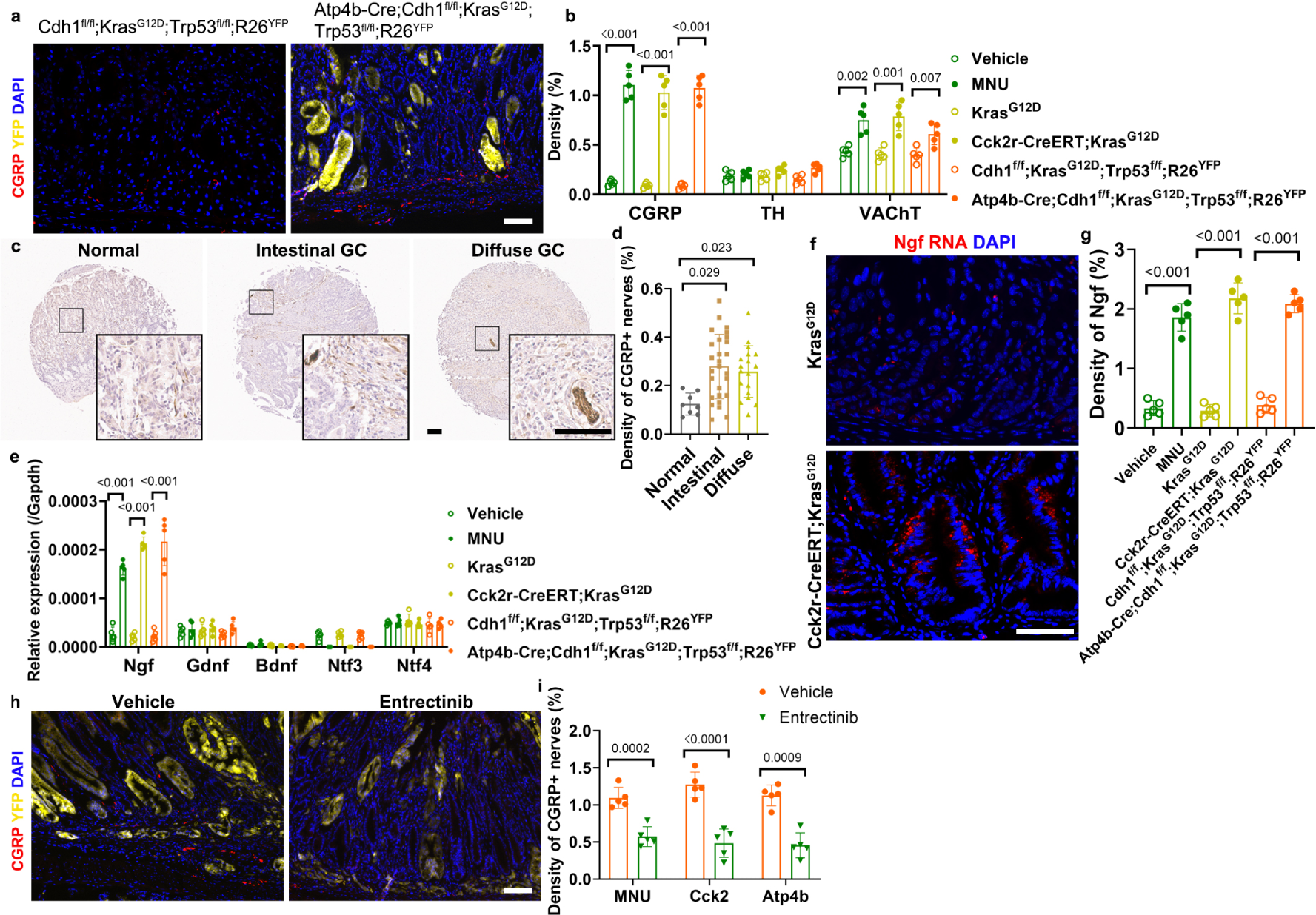
Sensory nerves are expanded in gastric cancers through NGF/TrkA signaling. (**a**) Representative images and (**b**) quantification of neural expansion in mouse gastric cancers (n = 5/ group). Scale bar, 100 μm. CGRP, Calcitonin gene-related peptide. TH, tyrosine Hydroxylase. VAChT, Vesicular acetylcholine transporter. (**c**) Representative images and (**d**) quantification of sensory nerve in human stomach and gastric cancer (n = 8 in normal group, 27 in intestinal GC group and 18 in diffuse GC group). Scale bar, 100 μm. (**e**) Relative expression of neurotrophin family in mouse gastric epithelial cells and mouse gastric cancer cells (n = 5/group). (**f**) Representative images and (**g**) quantification of Ngf in situ hybridization in mouse GCs (n = 5/ group). Scale bar, 100 μm. (**h**) Representative images and (**i**) quantification of sensory nerves in mouse gastric cancers treated by Entrectinib or Vehicle (n = 5/group). Scale bar, 100 μm. Data represent mean ± SEM, and *P* values were calculated by ANOVA in **d**, by t test in **b**, **e**, **g**, and **i**.

NGF is essential for the growth and survival of sensory neurons during development ^20^, and is frequently upregulated with inflammatory injury ^21^ ^22^ and cancer ^13^. Acute gastric injury following treatment of C57BL/6 mice with the protonophore DMP-777 or high dose MNU led to upregulation of NGF expression in gastric epithelial cells (**Extended Data Fig. 2a-b**). Expression of *Kras*^G12D^ mutation in gastric epithelial cells in *Mist1-*CreERT; *Kras*^G12D^; tGFP mice led to gastric dysplasia ^23^ and significantly increased NGF expression (**Extended Data Fig. 2c-d**). Analysis of neurotrophin gene expression in purified GC epithelial cells from multiple GC models showed NGF to be the most highly upregulated (**Fig. 1e**). In situ hybridization (RNAScope) confirmed that following Kras activation, NGF was significantly upregulated early on in the gastric isthmus region known to harbor gastric stem cells (**Fig. 1f-g and Extended Data Fig. 2e**). Overexpression of NGF throughout the gastric epithelium in the NGF knockin line (*Tff2*-Cre; *R26*-NGF) led to a 30-fold expansion of CGRP+ sensory nerves in the gastric mucosa **(Extended Data Fig. 2f)**. In addition, given that NGF has been shown to promote angiogenesis^24^, we analyzed microvessel density and confirmed it was increased in this line in close proximity to CGRP+ sensory nerves **(Extended Data Fig. 2g)**. The expansion of CGRP+ sensory nerves in GC mouse models was suppressed by the treatment with the Trk inhibitor Entrectinib (**Fig. 1h-i and Extended Data Fig. 2h**), indicating a dependence on NGF/Trk signaling.

### The stomach is innervated by nociceptive neurons and gastric cancer preferentially attracts CGRP+ peptidergic neurons

Retrograde tracing of the normal stomach was performed with three neural tracers (Fast Blue, CTB-Alexa Flur 594, and rAAV-hSyn-EGFP). One week after injection into the anterior or posterior gastric wall, the tracers were observed in neurons in the unilateral Nodose Ganglia (NG) and Dorsal Root Ganglia (DRG) T7 to T13, with only slight variation depending on the tracer (**Fig. 2a-b**). Immunostaining revealed that in NGs and DRGs, CGRP+ peptidergic nociceptive neurons and IB4+ nonpeptidergic nociceptive neurons were mutually exclusive subtypes (**Extended Data Fig. 3a**). CGRP+ peptidergic nociceptive neurons made up all of the retrogradely labeled stomach-innervating neurons in DRGs and 82.25% ± 4.99% of labeled neurons in NGs, with the few remainders being IB4+ nonpeptidergic nociceptive neurons (**Fig. 2c-d**). CGRP+ neurons expressed several other neuropeptides, including Substance P (SP) and somatostatin (SST) (**Extended Data Fig. 3b**). NGF-overexpression produced a large increase in CGRP+ nerves (**Extended Data Fig. 2f**), but no change in traced NGs and DRGs (**Extended Data Fig. 3c-e**), suggesting primarily axonogenesis.

**Figure 2.**
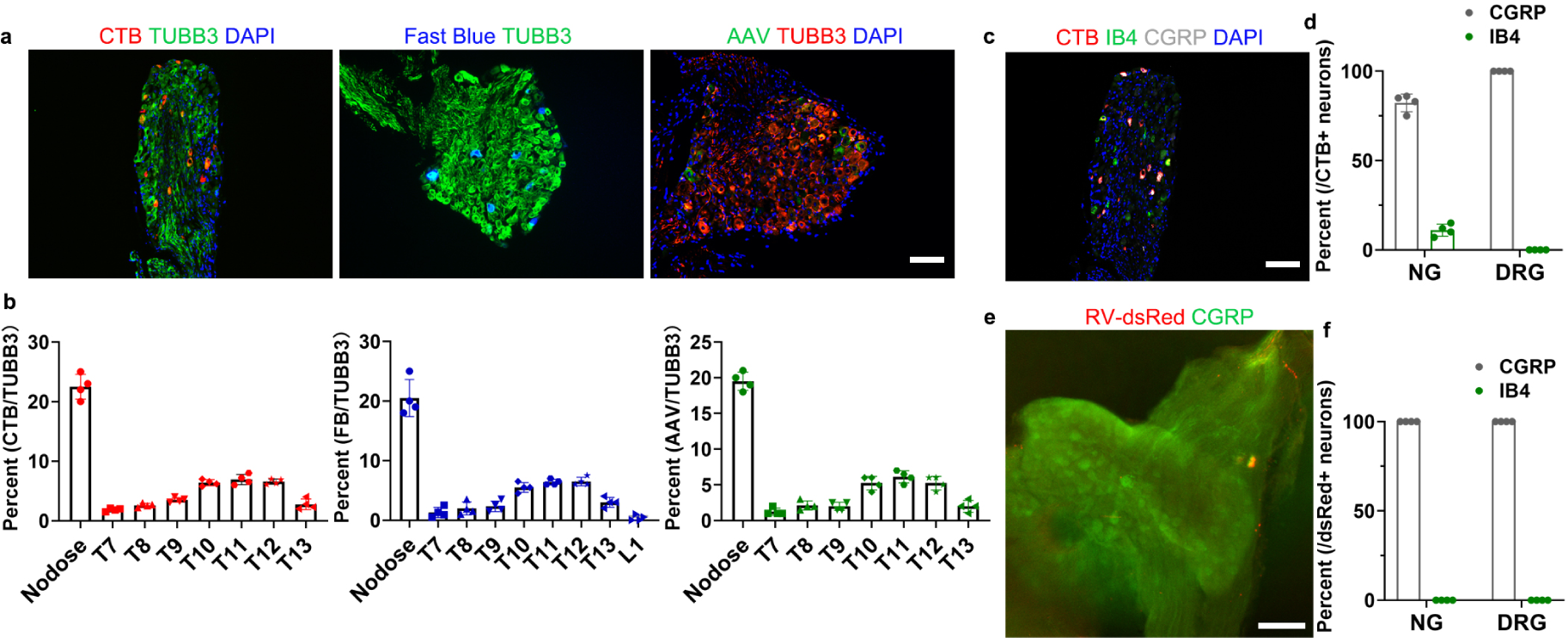
Identification of sensory neurons innervating to normal stomach and GC. (**a**) Representative images and (**b**) quantification of the stomach-innervating sensory neurons in nodose ganglia and dorsal root ganglia with three different neural tracers (n = 4/group). CTB, Cholera Toxin Subunit B-Alexa Flour 594. AAV, AAV-hSyn-eGFP. TUBB3, beta 3 tubulin. NG, nodose ganglia. DRG, dorsal root ganglia. Scale bar, 100 μm. (**c**) Representative images and (**d**) quantification of molecular subtypes of the stomach-innervating sensory neurons (n = 4/group). CGRP, calcitonin gene-related peptide. IB4, Isolectin B4. Scale bar, 100 μm. (**e**) Representative images of GC-innervating sensory neurons with whole mount staining and (**f**) quantification (n = 4/group). RV, Rabies virus. Scale bar, 100 μm. Data represent mean ± SEM.

Finally, we confirmed the innervation ACKP gastric orthotopic tumors using glycoprotein-deleted Rabies viruses (EnvA-ΔG-dsRed RV) ^27^. ACKP gastric cancer cells were infected with LV-EF1a-TVA-P2A-G lentivirus (Biohippo) to express TVA and glycoprotein (G). At 2 weeks post-injection, analysis revealed neurons labeled with dsRed detected in NGs and DRGs (**Fig. 2e**). Notably, all of the labeled neurons expressed CGRP (**Fig. 2e-f**), indicating that GC cells preferentially attract CGRP+ peptidergic nociceptive neurons.

### Sensory neuropeptide CGRP and its receptor Ramp1 are upregulated in GC

Consistent with CGRP+ nerve expansion, we found significantly higher levels of CGRP mRNA in stomach-innervating ganglia (**Fig. 3a**) and CGRP peptide in GC mouse stomachs (**Fig. 3b**). Similar increases in CGRP mRNA and CGRP peptide were found in the ganglia and stomachs, respectively, of *Tff2*-Cre; *R26*-NGF mice (**Extended Data Fig. 4a-b**), consistent with known effects of NGF on CGRP ^29^. Treatment of GC mice with the Trk receptor antagonist Entrectinib resulted in reduced CGRP synthesis and release (**Fig. 3a**). The CGRP receptor consists of a heterodimer, comprising the calcitonin receptor-like receptor (CRL) encoded by *CALCRL* and the receptor activity modifying protein 1 (RAMP1) ^30^. Analysis of available single-cell transcriptome data from mouse (GSE157694 and GSE116514) and human stomach (OMIX001073) showed that both Calcrl and Ramp1 were expressed in normal gastric epithelial and stromal cells (**Fig. 3c-d**), while the expression levels of other neuropeptide receptors were much lower (**Extended Data Fig. 4c)**. There was a marked increase in Ramp1 expression in GC cells (and fibroblasts) (**Fig. 3d**), with some Ramp1+ cells showing strong Ki-67 expression (**Extended Data Fig. 4d**) indicating active proliferation.

**Figure 3.**
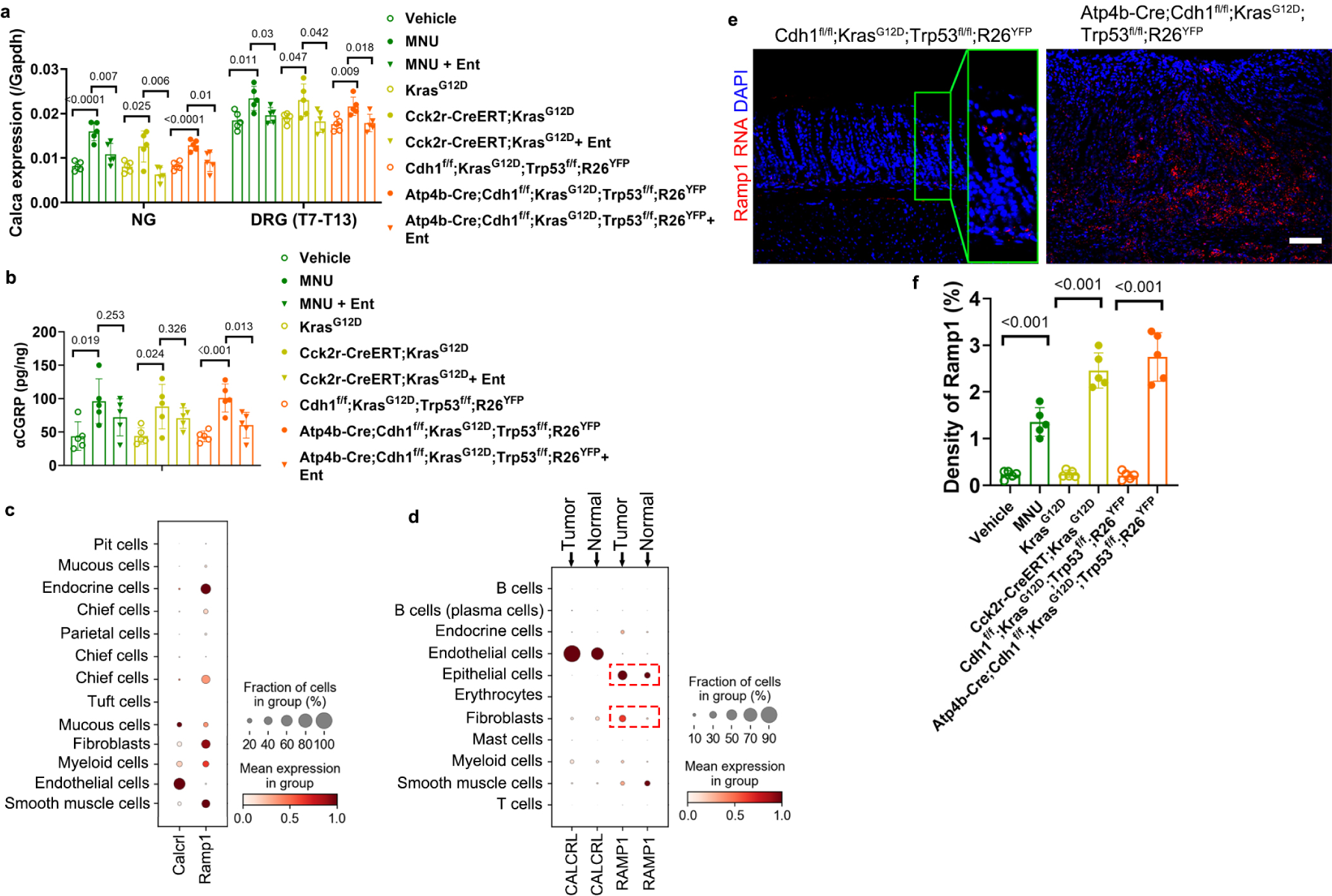
Sensory neuropeptide CGRP and its receptor Ramp1 are aberrantly elevated in GC. (**a**) Expression levels of *Calca*, a gene encoding for CGRP, in the stomach-innervating and GC-innervating sensory neurons (n = 5/group). Ent, Entrectinib. (**b**) Concentrations of CGRP peptide in the stomach and GC (n = 5/group). (**c**) Dot plot of single-cell transcriptome data from mouse stomach (GSE157694 and GSE116514). (**d**) Dot plot of single-cell transcriptome data from human stomach and GC (OMIX001073). (**e**) Representative images and (**f**) quantification of in situ hybridization (RNAScope) of Ramp1 RNA in mouse stomach and gastric cancers (n = 5/group). Scale bar, 100 μm. Data represent mean ± SEM, and *P* values were calculated by ANOVA in **a** and **b**, by t test in **f**.

In situ hybridization/RNAScope of the normal stomach revealed that Ramp1 was expressed in the isthmus/stem cell region of the gastric glands, as well as in chief cells (**Fig. 3e and Extended Data Fig. 4e**). Ramp1 was highly expressed in most malignant cells and Ramp1 staining in GC mice was much higher than that in controls (**Fig. 3e-f**), while the density of Calcrl staining in GC mice was similar to controls (**Extended Data Fig. 4f-g**). Analysis of available bulk RNA sequencing data (OMIX001073) revealed significant upregulation of RAMP1 in human GCs, with high expression of RAMP1 correlating with poor overall survival (**Extended Data Fig. 4h-i**).

### Nociceptive neurons promote gastric hyperplasia and GC development through CGRP signaling

We utilized a model of chemogenetic activation (*R26*-hM3Dq) in which the DREADD agonist clozapine N-oxide (CNO) is used to induce CGRP release from nociceptive neurons ^18^. As *Calca*-Cre;*R26*-hM3Dq mice did not tolerate well chronic activation, we crossed *R26*-hM3Dq mice with *Trpv1*-Cre, as Trpv1 nociceptive neurons show strong overlap with CGRP+ nociceptive neurons ^32^. *Trpv1*-Cre;*R26*-hM3Dq mice were treated with CNO given on alternate weeks for a total of 40 weeks, which led to marked gastric hyperplasia with increased proliferation (**Fig. 4a-c**). Further, activation in *Trpv1*-Cre;*R26*-hM3Dq mice of nociceptive neurons with CNO for 10 days in a gastric ulcer model^34^ also led to faster ulcer healing after injury, with higher Ki-67 expression and reduced ulcer size (**Extended Data Fig. 5a-c**). Notably, activation of nociceptive neurons in *Trpv1*-Cre;*R26*-hM3Dq mice significantly accelerated MNU-induced tumor growth **(Fig. 4d-e**). CNO-activated mice treated with a diet containing Rimegepant (a CGRP antagonist) showed reduced MNU-induced tumor development compared to untreated controls (**Fig. 4d-e**). In the syngeneic orthotopic tumor allograft model, nociceptive neuronal activation resulted in significantly increased tumor volume by MRI imaging and decreased survival, which was again inhibited by Rimegepant treatment (**Fig. 4f-h and Extended Data Fig. 5d**).

**Figure 4.**
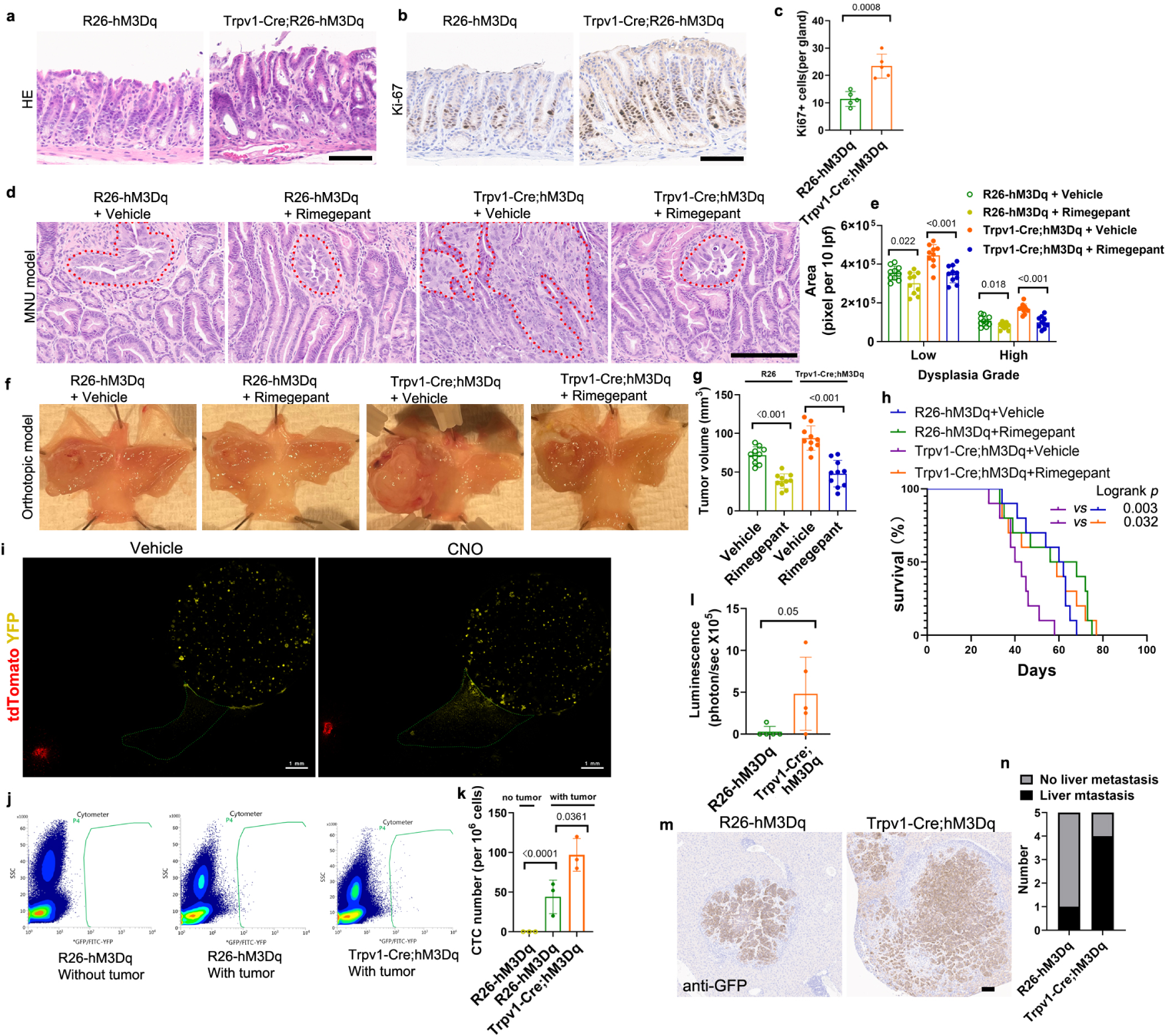
Nociceptive neurons promote GC development and metastasis through CGRP signaling. (**a**) Representative images of HE staining in the stomach from CNO-treated mice (n = 5/group). Scale bar, 100 μm. (**b**) Representative images and (**c**) quantification of Ki-67 staining in the stomach from CNO-treated mice (n = 5/group). Scale bar, 100 μm. (**d**) Representative images and (**e**) quantification of MNU-induced dysplasia from CNO-treated mice with or without Rimegepant administration (n = 10/group). Scale bar, 100 μm. (**f**) Representative images of syngeneic orthotopic tumors from CNO-treated mice with or without Rimegepant administration (n = 10/group). (**g**) Quantification of tumor volume calculated from MRI. (**h**) Kaplan-Meier curve of survival data from (**f**). (**i**) DRG (red) from *Trpv1*-Cre; hM3Dq; tdTomato mouse was cocultured with gastric cancer spheroids (yellow) and was treated with CNO. Representative images showed cancer cell migrating towards DRG. Scale bar, 1 mm. (**j**) Representative images and (**k**) quantification of circulating tumor cells in the orthotopic model. (**l**) quantification of luminescence from spontaneous metastatic mice (n = 5/group). (**m**) Representative images of liver metastasis samples and cancer cell staining (GFP). Scale bar, 100 μm. (**n**) Quantification of liver metastasis rate. Data represent mean ± SEM, and *P* values were calculated by ANOVA in **k**, by t test in **c**, **e**, **g**, and **l**.

To test whether nociceptive nerves are necessary for GC progression, we ablated nociceptive neurons in DRGs and NGs ^19^ in *Trpv1*-Cre;*R26*-iDTR mice using diphtheria toxin (DT) injections. DT-treated mice showed a significant decrease in nociceptive neurons (**Extended Data Fig. 5e-f**). DT treatment suppressed MNU-induced tumor development in *Trpv1*-Cre;*R26*-iDTR mice (**Extended Data Fig. 5g**), and inhibited growth of orthotopic allograft tumors (**Extended Data Fig. 5h**).

CGRP increased the spheroid-forming capacity of ACKP cells in vitro in a 3D Matrigel culture system. Treatment with murine CGRP peptide significantly increased the size of spheroids (**Extended Data Fig. 5i-j**) and cell proliferation as quantified by Edu staining (**Extended Data Fig. 5k-l**), which was blocked by Rimegepant treatment. A cell viability assay showed that CGRP promoted the growth of GC spheroids in a concentration-dependent manner (**Extended Data Fig. 5m**). Interestingly, activation of nociceptive nerves with CNO also led to increased Pdgfra+ CAFs in gastric orthotopic tumors (**Extended Data Fig. 6a-b**). CGRP sustained the proliferation of gastric CAFs in a concentration-dependent manner (**Extended Data Fig. 6c-d**), with CGRP treatment leading to a significant (8-fold) upregulation in CAFs of Il-6 (**Extended Data Fig. 6e**). Thus, CGRP+ nociceptive nerves appear to promote GC development through effects on GC cells and CAFs.

### Nociceptive neurons promote liver and lymph node metastasis

Nociceptive neurons have been reported to contribute to cancer metastasis ^35^. In a coculture model, CNO-dependent activation of nociceptive neurons in DRG significantly promoted the migration of cancer cells towards the DRG (**Fig. 4i**). In the ACKP orthotopic allograft model, CNO-dependent nociceptive nerve activation resulted in a significant increase of circulating tumor cells (CTCs) in the portal vein (**Fig. 4j-k**). We established a spontaneous gastric metastatic model through esophagojejunostomy with Roux-en-Y anastomosis performed 4 weeks post-orthotopic implantation of luciferase-expressing ACKP cells (**Extended Data Fig. 7a**). Following primary tumor resection, liver metastases were detectable by bioluminescence from the third week (**Extended Data Fig. 7b**). CNO-treated *Trpv1*-Cre;*R26*-hM3Dq mice had higher bioluminescence reflecting increased liver metastases that was pathologically confirmed (**Fig. 4l-m**). Four out of five *Trpv1*-Cre;*R26*-hM3Dq mice showed liver metastases and 3 had lymph node metastases, while only one mouse in the control group had liver metastases with no lymph node metastases (**Fig. 4n**). Liver mets in *Trpv1*-Cre;*R26*-hM3Dq mice showed higher expression levels of Ki-67 and a higher density of CGRP+ nociceptive nerves (**Extended Data Fig. 7c-f**), while, CGRP+ nerves in normal or tumor-adjacent liver were nearly undetectable (**Extended Data Fig. 7e-f**). CGRP+ nerves were also found in lymph nodes with mets (**Extended Data Fig. 7g**). Liver metastatic tumors in *Trpv1*-Cre;*R26*-hM3Dq mice had more α-SMA+ activated hepatic stellate cells (HSCs) than controls or tumor adjacent liver (**Extended Data Fig. 7h-i**) and analysis of available single-cell transcriptome data from mouse liver ^36^ showed that the CGRP receptor, Ramp1, was highly expressed in HSCs (**Extended Data Fig. 7j**). These data suggest that nociceptive neurons also promote the growth of liver metastatic GC tumors.

### Nociceptive neurons directly interact with gastric cancer cells

CGRP released from afferent nerves can exert paracrine effects on surrounding tissue ^37^, but direct interactions with between sensory nerves have not been shown for extra-cranial tumors. We observed that nociceptive fibers were closely juxtaposed with GC cells (**Fig. 1**), and thus placed in a 3D coculture model both DRGs from *Trpv1*-Cre; hM3Dq; tdTomato mice and GC spheroids. The nociceptive neurons generated synapse-like structures on the GC spheroids (**Fig. 5a**), and with activation by CNO treatment, GC cells migrated along the nerves towards the DRG in a manner that could be blocked by Rimegepant treatment (**Fig. 5a**). GC spheroids that exhibited neuronal connections were analyzed, which revealed expression of a number of chemical synaptic genes, some of which were upregulated in neuron-connected cancer cells (**Extended Data Fig. 8a**). In particular, Neurod1 and Ascl1^38^, two key transcription factors known to be involved in neuronal reprogramming, were upregulated, suggesting that cancer cell-neuron connections may induce a neuronal-like phenotype to promote migration (**Fig. 5a),** analogous to effects with neuro-gliomal synapses ^5^.

**Figure 5.**
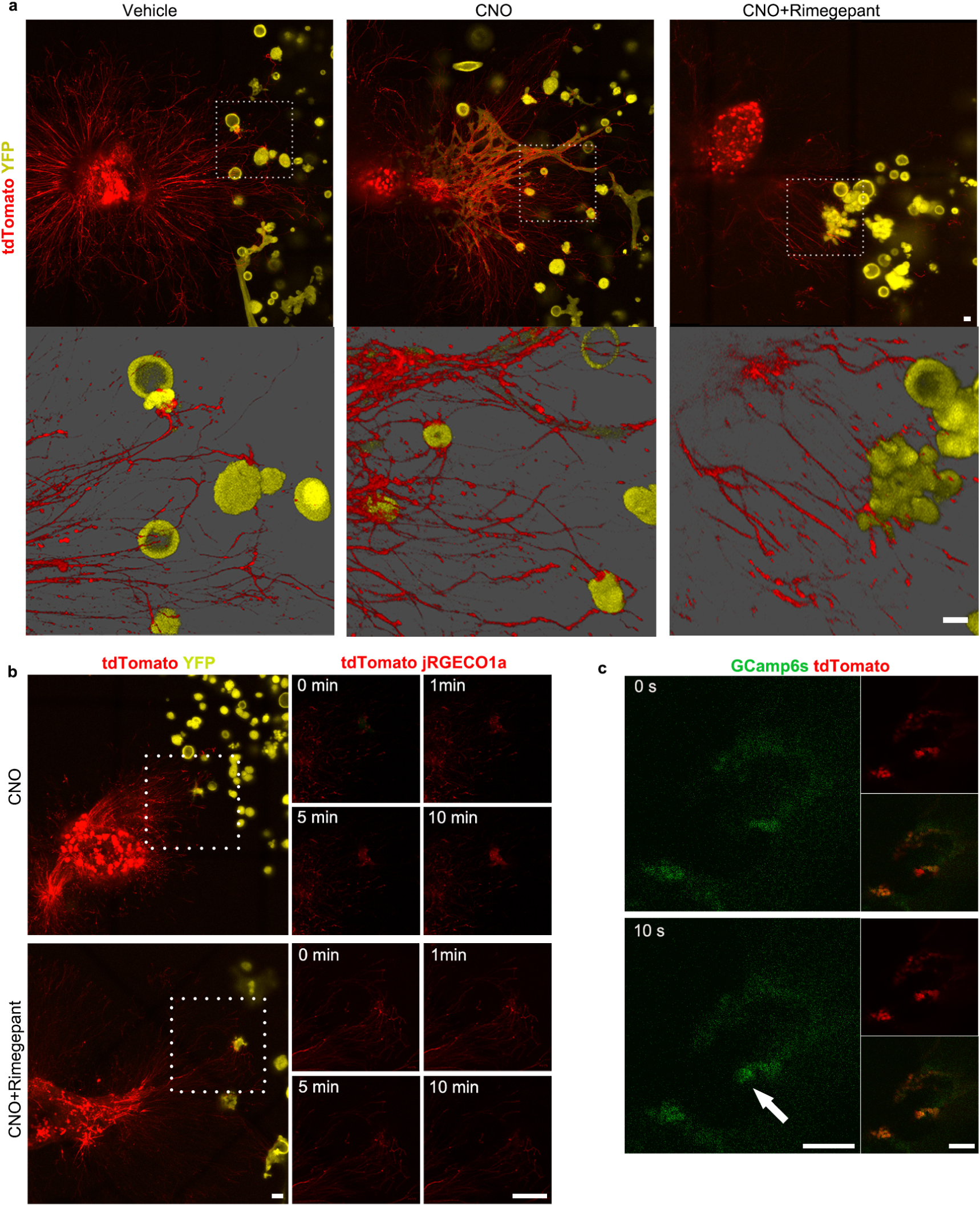
Nociceptive neurons directly interact with gastric cancer cells. (**a**) Representative images of proximity coculture of DRG (from *Trpv1*-Cre; hM3Dq; tdTomato mouse) and GC spheroids. High magnification 3D images showed synapse-like structures. Scale bar, 100 μm. (**b**) Once proximity coculture of DRG (from *Trpv1*-Cre; hM3Dq; tdTomato mouse) and GC spheroids (infected with rAAV-CMV-jRGECO1a) formed synapse-like structures, nociceptive neurons were treated with CNO or CNO+Rimegepant and calcium indicator jRGECO1a (red) in GC spheroids was monitored. Scale bar, 100 μm. (**c**) Representative images of in vivo calcium imaging in nodose ganglia of *Trpv1*-Cre; GCamp6s; tdTomato mice. ACKP-ChR2 tumor was optogenetic activated by 473 nm blue light. The white arrow showed activation of nociceptive neurons. Scale bar, 100 μm.

Nociceptive nerves are known to release neuropeptides and also sense molecules such as glutamate, ATP, NGF, etc.^39^., and we hypothesized that the membrane protrusions might represent synapse-like structures. Gastric cancer spheroids were infected with rAAV-CMV-NES-jRGECO1a, placed in 3D coculture with DRGs from *Trpv1*-Cre; hM3Dq; tdTomato mice and analyzed with calcium imaging. Activating peptidergic nociceptive neurons with CNO induced slow calcium flux in neuron-connected cancer cells, which was blocked by Rimegepant treatment (**Fig. 5b**), demonstrating direct signaling from CGRP+ neurons. The slower time frame suggests calcium release from endoplasmic reticulum rather than through voltage-gated calcium channels, and Ramp1 has been shown to induce calcium release through PLC-β1 ^29^. We then implanted ACKP GC cells infected with pLV-ChR2 lentivirus expressing Channel Rhodopsin in the anterior stomach wall of *Trpv1*-Cre;GCamp6s;tdTomato mice. Two weeks later, optogenetic activation and depolarization of ACKP GC cells led to fast calcium flux of nociceptive neurons in the ipsilateral NG as revealed by in vivo calcium imaging (**Fig. 5C, Supplementary video 1**). Moreover, optogenetic activation of ACKP cells resulted in increased CGRP peptide levels in tumors (**Extended Data Fig. 8b**), reflecting CGRP release from activated nociceptive neurons. Together, these data define a cancer cell-peptidergic neuronal circuit.

### Nociceptive neurons regulate gastric cancer through Rb/E2F pathway

Principle component analysis (PCA) on bulk RNA sequence data from GC spheroids alone, cultured alone or cocultured with either DRG or CNO-activated DRG, showed that unconnected spheroids clustered independently from neuron-connected spheroids cocultured with DRG or CNO-activated DRG (**Extended Data Fig. 8c**). Interestingly, gene set enrichment analysis (GSEA) indicated the Rb/E2F pathway to be significantly enriched in every comparison (**Extended Data Fig. 8d**). Differential expression analysis revealed that proliferation genes and Rb/E2F inducers were upregulated with Rb/E2F suppressors decreased (**Extended Data Fig. 8e**), pointing to a role for the Rb/E2F pathway in nociceptive neuron-dependent gastric cancer progression.

While binding of CGRP to Calcrl/Ramp1 leads to activation of multiple signaling pathways^29,43^, we combined CGRP treatment with different kinase inhibitors and assessed E2F activity. CGRP treatment alone in GC cells significantly increased E2F activity (**Extended Data Fig. 8f**), but among four kinase inhibitors, Wortmannin and KN-93 significantly suppressed CGRP-dependent E2F activity (**Extended Data Fig. 8f**), implicating the PI3K/Akt and CaMK pathways, respectively in E2F signaling by CGRP. Phosphorylation of Rb inhibits E2F suppressor activity and we confirmed increased p-Rb expression in orthotopic ACKP cells in CNO-activated *Trpv1*-Cre;hM3Dq; mice (**Extended Data Fig. 8g-h)**. Staining of 50 human GC samples with p-Rb antibody demonstrated that p-Rb was moderately expressed in 20% of GC patients and strongly expressed in 18% patients, with a significant correlation between CGRP immunoreactivity and p-Rb immunoreactivity (**Extended Data Fig. 8i**), supporting a role for the CGRP/Rb/E2F pathway in human GC.

## Discussion

We find in this study that, similar to gliomas, solid extra-cranial tumors such as gastric adenocarcinoma also develop within a complex neural circuitry. Distinct from gliomas, this involves predominantly peptidergic sensory nerves. Sensory nerves are attracted to GC largely through an NGF/Trk pathway, and activation of these sensory nerves promotes the growth and metastatic spread of GC via a CGRP-RAMP1 axis. In contrast to gliomas, the interaction between tumors and nerves does not involve excitatory electrochemical synapses but is mediated instead through slower paracrine signaling. Nevertheless, evidence for such a neural circuit is supported by the finding that in vivo optogenetic activation of gastric cancer cells leads to signaling in TRPV1 expressing Nodose Ganglia cells. In addition, stimulation of sensory nerves by GC cells leads to increased secretion of CGRP peptide consistent with a feed-forward loop which is proposed in this study (**Extended Data Fig. 9**). Similarly, chemogenetic activation of TRPV1 sensory nerves results in calcium signaling in GC cells, which could be blocked with a CGRP inhibitor. Taken together, these data indicate that GC co-opt the existing sensory nervous system to establish a gastric cancer-sensory nerve circuit that can drive gastric cancer progression.

Nerves are increasingly recognized as an integral part of the tumor microenvironment but their direct interactions with cancer cells have been most thoroughly elucidated in studies of CNS tumors^2,3,44^. GC was shown to be dependent on cholinergic innervation, as surgical vagotomy led to marked decrease in tumor growth, due in part to reduced muscarinic signaling ^11^. However, studies elsewhere have demonstrated an important role of sensory nerves in the growth of basal cell carcinomas ^45^, pancreatic cancer ^46^ and triple-negative breast cancer ^35^, and in fact sensory nerves are a component of vagal axons ^12^. The studies here suggest a much greater expansion of sensory nerves in response to tumor-derived signals such as NGF, and the findings that sensory nerve ablation abrogates tumor growth indicate the dominant role that sensory nerves play in modulating tumor growth.

The innervation of the stomach was elucidated through retrograde tracing using three different tracers, which showed neuronal innervation arising in the Nodose Ganglia (NG) and in Dorsal Root Ganglia (DRG, T7-T13). Overall, peptidergic nociceptive neurons expressing CGRP were dominant, in contrast to an earlier study^47^, and were markedly increased by NGF in gastric cancer. The interactions between CGRP+ sensory nerves and GC cells appeared to be direct and could be visualized in spheroid culture as synapse-like structure. Optogenetic activation showed that GC cells signaled directly to sensory nerves. Since membrane depolarization has been proposed as a biomarker for cancer ^40,41^, we posit that depolarization-evoked exocytosis ^42^ in cancer cells may lead to release of small molecules such as ATP, glutamate, and hydrogen ion, which can then bind to voltage-gated ion channel receptors on sensory neurons. GC cells also receive signals from nerves, but the calcium responses to chemogenetic/CGRP stimulation was slower than that in a true synapse and likely mediated by internal calcium release rather than from voltage gated channels. However, GC cells with neural connections showed upregulation of two transcription factors involved in neuronal reprogramming (Neurod1 and Ascl1), which may facilitate communication and contribute to cancer spread.

Sensory nerves can interact directly with epithelial cells in the normal gut ^16–19^, with regulation for example of enteroendocrine cells ^16^, and of goblet cells ^18^ to drive mucous production. However, we show here that gastric expression of the CGRP receptor RAMP1 could be seen not only in chief cells but also in gastric isthmus region where stem/progenitor cells have been localized. Chemogenetic activation of sensory nerves led to gastric hyperplasia, consistent with effects on normal gastric epithelial progenitors, and CGRP treatment of gastric cancer organoids, where RAMP1 is upregulated, led to increased spheroid growth and survival. CGRP-expressing sensory nerves have previously been shown to act directly on haematopoietic stem cells to increase their mobilization ^48^, and thus we speculate that gastric stem cells may be similarly regulated, but here through a PI3K/Akt and CaMK pathways leading to activation of the Rb/E2F pathway. However, we also found in this study that CGRP+ sensory nerves likely interacted with stromal cells which expressed RAMP1 and CALCR. RAMP1 was highly expressed in hepatic stellate cells, and upregulated in gastric CAFs, which showed increased proliferation and IL-6 secretion in response to CGRP treatment. Indeed, CGRP has been shown to regulate many aspects of tumor biology, including promoting cancer cell cytoprotective autophagy ^9^, increasing the exhaustion of cytotoxic T cells ^8^, and enhancing tumor growth ^49^ in a paracrine manner. The findings from our murine models were supported in our human data sets, as CGRP+ sensory nerves were markedly expanded in tissue microarrays of human gastric cancer samples. Furthermore, the main downstream target of CGRP signaling in cancer, phosphorylated Rb (p-Rb) was expressed in over a third of patients and correlated with CGRP expression. Antagonism of CGRP receptors with Rimegepant, an FDA approved drug, consistently reduced GC growth in vitro and in vivo in all GC models studied, prolonging survival. In migraine patients, this class of drugs is generally well tolerated with minimal adverse effects ^50^. Interestingly, a population-based cohort study in Denmark ^51^ showed that the incidences of gastrointestinal cancers (especially gastric cancer) were significantly lower in migraine patients. In summary, we demonstrate a cancer cell-nociceptive neuronal circuit that critically promotes GC development. Our findings suggest that anti-CGRP represents a promising preventive and therapeutic strategy for GC.

## Author contribution

XZ and TCW conceived, designed the study and wrote the manuscript. XZ, FW, JQ, YO and GL performed most of the experiments. EM performed the computational analyses of scRNA sequencing data. SR generated and characterized Tcon mice and ACKP cells. EM, DC and SR participated in the discussion. TCW generated the concepts, analyzed and interpretate the data, developed the idea and hypothesis, secured the funding.

## Acknowledgements

This work was supported by grants UO1DK103155, R01DK128195, R01CA272901, R01CA224428, W81XWH-21-10901 as well as an NCI Outstanding Investigator Award (R35CA210088) to TCW. SR is supported by R01CA272891 and P50CA127003 grants. This research was supported in part through the NIH/NCI Cancer Center Support Grant P30CA013696 and used the resources of the Herbert Irving Comprehensive Cancer Center Flow Cytometry Shared Resources, Molecular Pathology (MPSR), Genomics and High Throughput Screening, the Oncology Precision Therapeutics and Imaging Core (OPTIC) and the Genetically Modified Mouse Model Shared Resource (GMMMSR). This research was also supported by the Columbia University Digestive and Liver Disease Research Center (CU-DLDRC) P30DK132710 grant and its Bio-Imaging, Organoid, and Bioinformatics/Single-Cell Analysis Cores. We thank the core directors, managers, and staff for their expert assistance with our studies. We are also grateful for support from Tonix Pharmaceuticals and the Torrey Coast Foundation. We acknowledge the support of the Core facilities, especially the Animal Facility at the Institute of Comparative Medicine at Columbia University. We thank the members of the Wang and Ryeom laboratories for helpful input throughout the course of this study.

## Methods

### EXPERIMENTAL MODEL AND SUBJECT DETAILS

#### Animals

All animal experiments were conducted in accordance with the National Institute of Health guidelines for animal research and approved by the Institutional Animal Care and Use Committee of the Columbia University. Mice were housed in a specific pathogen-free facility. *Tff2*-Cre ^13^, *R26*-NGF ^13^, *Cck2r*-CreERT ^14^, and LSL-*Kras*^G12D^ ^52^ were described previously. *Trpv1*-Cre, *Calca*-Cre, *R26*-hM3Dq, *R26*-iDTR, and *R26*-tdTomato were purchased from the Jackson Laboratory. Tcon mice were generated as previously ^15^.

#### Mouse gastric cancer models

MNU-induced gastric cancer model was generated as previously ^14^. Briefly, age and sex matched 8 to 12-week old C57BL/6 mice were given drinking water containing 240 ppm MNU on alternate weeks for a total of 10 weeks. Cancer samples were collected at 10 months. *Cck2r*-CreERT; LSL-*Kras*^G12D^ mice were administered 2 mg tamoxifen in 200 μL corn oil by gavage. Cancer samples were collected at 8 months. Tcon mice were generated by cross breeding *Atp4b*-Cre transgenic mice with *Cdh1*^fl/fl^, LSL-*Kras*^G12D^, *Trp53* ^fl/fl^, and *R26*-YFP mice ^15^. Cancer samples were collected at 10 weeks. For syngeneically orthotopic allografts, ACKP cells ^15^ (gastric cancer cells derived from *Atp4b*-Cre; *Cdh1*^fl/fl^; *Kras*^G12D^; *Trp53* ^fl/fl^; YFP mice) were used. Mice were anaesthetized with 2–4% isoflurane and placed on a heat pad. A middle incision was performed, and the anterior wall of stomach was exposed. ACKP cells (1×10^6^ cells in 10 μL PBS) were drawn up into a sterile 31 G × 5/8” syringe. These cells were injected slowly into the middle of the anterior wall without dissemination. Cancer samples were collected at 4 weeks.

#### Cell culture

Murine gastric cancer cells (ACKP cells) derived from *Atp4b*-Cre; *Cdh1*^fl/fl^; *Kras*^G12D^; *Trp53* ^fl/fl^; YFP mice were cultured in DMEM medium supplemented with 10% FBS and 1% penicillin/streptomycin in a humidified incubator at 37 ℃, as described previously ^15^.

#### Human samples

Human tissue arrays containing normal stomach samples (ST1001) and gastric cancer samples (ST1003b) were purchased from Amsbio.

### METHOD DETAILS

#### Preparation of reagents

MNU (Gojira Fine Chemicals) at high dose (2400 ppm) for acute gastric injury was administered by oral gavage. MNU at low dose (240 ppm) for gastric tumorigenesis was dissolved in drinking water and freshly prepared twice per week. DMP-777 (MedChem Express) was dissolved in dimethyl sulfoxide and administrated orally as a gavage (250 mg/Kg) once daily. Murine CGRP peptide (Tocris) was dissolved in PBS and given at a concentration of 1 μM in vitro. Entrectinib (Selleckchem) and Rimegepant (MedChem Express) were mixed into the AIN-76A mouse chow (100 mg/Kg for Entrectinib; 10 mg/Kg for Rimegepant) and fed for the indicated periods. For chronic activation of Designer Receptors Exclusively Activated by Designer Drugs (DREADD)-controlled neuronal activity, the ligand clozapine-N-oxide (CNO) was administered through drinking water at a concentration of 4 μg CNO/ mL ^33^.

#### Retrograde tracing

The mice were anesthetized by isoflurane. A laparotomy was performed to expose the stomach. The greater curvature of the stomach was held with microsurgical forceps. Cholera toxin subunit B conjugated with Alexa Fluor 594 (0.5%, Invitrogen), Fast Blue (5%, Polysciences) or rAAV-hSyn-EGFP (10^12^vp/ml, eBioHippo) was injected slowly at several spots beneath the serosa of the whole anterior wall or posterior wall with a Hamilton syringe (30G, 25 μL). The needle was kept in place for 10 seconds to avoid dye leakage. The injection site was rinsed with saline to remove any leaking dye. Mice were sacrificed ten days after injection. Nodose ganglia and dorsal root ganglia (T3-L3) were dissected.

#### Immunohistochemistry and immunofluorescence

Tissues were fixed overnight in 4% paraformaldehyde or 10% formalin at 4 ℃, embedded into paraffin block or OCT compounds. The slides were processed by standard histological methods. Following antigen retrieval, the tissues were blocked with 10% bovine serum albumin (BSA) (Roche) for 1 h at room temperature, and then stained with primary antibodies at 4 ℃ overnight. The following primary antibodies were used: anti-CGRP (1:200, Abcam, Sigma-Aldrich), anti-VAChT (1:200, Synaptic Systems), anti-TH (1:200, Millipore), anti-E-cadherin (1:200, Cell Signaling Technology), anti-Cd31 (1:200, R&D Systems), anti-Tubb3 (1:200, Abcam), IB4 (1:200, Vector Laboratories), anti-Ki67 (1:50, Abcam), anti-GFP (1:200, Abcam), anti-Phospho-Rb (Ser807/811, 1:200, Cell Signaling Technology), anti-NF200 (1:200, Novus Biologicals), anti-Substance P (1:200, Abcam), anti-Somatostatin (1:200, Millipore), anti-Cd8a (1:200, Abcam), anti-F4/80 (1:200, Cell Signaling Technology), anti-Pdgfra (1:200, Cell Signaling Technology), anti-α-SMA (1:500, Abcam). The sections were incubated with biotinylated secondary antibodies (1:200, Vector Laboratories) for 1 h, and visualized with a standard avidin biotinylated peroxidase complex method. For immunofluorescence, Alexa Fluor 488, 594 or 647 secondary antibodies (Thermo Fisher Scientific) were used. EdU staining was done following the manual instruction of the kit (Thermo Fisher Scientific).

#### RNAscope in situ hybridization

RNAscope in situ hybridization (ISH) was performed with RNAscope Multiplex Fluorescent Reagent Kit v2 (Advanced Cell Diagnostics). The slides were baked in a dry oven for 1 h at 60 ℃, followed by deparaffinization. The tissues were then treated with hydrogen peroxide. When hydrophobic boundaries were created, protease reagent was added to each section. Channel 1 or Channel 2 probes were pipetted onto each section until fully submerged. Following probe incubation, the slides were washed two times in RNAscope wash buffer and returned to the oven for 30 minutes after submersion in AMP reagent. Channel 1 (Opal 570) was for Ngf and Channel 2 (Opal 570) was for Ramp1. Slides were then covered with antifade mounting medium with DAPI.

#### Quantitative analysis of mRNA expression

Total RNA was extracted from the tissue and cells with TRIzol reagent (Invitrogen) and was reverse transcribed into cDNA with qScript cDNA SuperMix (QuantaBio). Expression levels of indicated genes were quantified by qPCR assays using a 7500 Real-time PCR system (Applied Biosystems) with a SYBR Green FastMix kit (QuantaBio). Predesigned sequences from Integrated DNA Technologies were used in this experiment and available upon request.

#### Bulk RNA sequencing

Gastric cancer spheroids were cocultured with DRG from Trpv1-Cre;hM3Dq;tdTomato mouse. We treated the coculture with CNO (10 μM) to activate sensory neurons for five days. RNA was isolated from neuron-connected spheroids cocultured with DRG, neuron-connected spheroids cocultured with CNO-activated DRG, and spheroids alone using the RNeasy Mini Kit (QIAGEN). RNA quality was confirmed using the Advanced Analytical Fragment Analyzer. 100 bp paired-end reads were sequenced on the Illumina HiSeq 2500 system to at least 40 million reads per sample at the Genome Center Facility at Columbia University. Differentially expressed genes were determined using DESeq2 version 1.24.0. Gene set enrichment analysis was performed using GSEA 4.3.2.

#### Enzyme linked immunosorbent assay

ELISA for murine CGRP (MyBioSource) was performed according to manufacture’s protocols. The standard curves were calculated by Curve Expert. Stomach samples were collected and washed by PBS to remove blood. The samples were then put into cold PBS (250 μL for 1 stomach) containing a protease/phosphatase cocktail. The samples were fully lysed and centrifuged at 6000 rpm for 15min. The supernatant was extracted for ELISA assay.

#### Magnetic resonance imaging

The mice were anesthetized by isoflurane. MRI was done using a Bruker Biospec 9.4T Small Animal MR Imager at Oncology Precision Therapeutics and Imaging Core at Columbia University. The volume of gastric cancer was calculated with ImageJ.

#### Flow cytometry

Epithelial cells and stromal cells were isolated from the mouse stomach as previously described ^14^. Briefly, the stomach or gastric tumor samples were collected, flushed with cold PBS. Then, the tissues were minced finely using scissors and placed in 20 mL of digestion media (HBSS, 1% HEPES, 1 mg/mL Collagenase P, 1 mg/mL Dispase II, 10 μg/mL DNAse I, and 1% BSA) and incubated at 37 ℃ for 20 minutes on a rotator. The solution was filtered through a 40 μm filter and then placed in 5 mL RBC lysis buffer (BioLegend) for 5 minutes. The reaction was stopped by PBS. Before the cells were stained with antibodies, they were incubated with TruStain FcX (BioLegend) for 5 minutes on ice. Finally, the suspension was stained with the following antibodies: EpCAM-APC (1:200, BioLegend), CD45-BV785 (1:200, BioLegend), CD45-BV421 (1:200, BioLegend), CD11b-BV650 (1:200, BioLegend), F4/80-BUV496 (1:200, BioLegend), CD206-BV711 (1:200, BioLegend), CD163-PE-Cy7 (1:200, BioLegend), CD31-PE (1:200, BioLegend). DAPI (BioLegend) or Zombie NIR (BioLegend) was used as a viability dye. To analyze the circulating tumor cells, blood was harvested from portal vein. CTCs were defined as YFP+ cells.

#### Spontaneous metastasis model

Liver metastasis can be found within a week by portal vein injection of cancer cells ^53^. However, due to the fast progression of liver metastasis in that model, there might be no time for tumor innervation. Here, we established a spontaneous metastasis model from gastric orthotopic allograft. Firstly, gastric orthotopic allografts were done as described previously. Four weeks later, we resected the stomach as well as the tumor followed by esophagojejunostomy with Roux-en-Y anastomosis ^54^. Then the mice were closely monitored and sacrificed after four more weeks.

#### 3D#cancer spheroid culture

ACKP cells were resuspended in Matrigel (1:1, Corning), and 5000 cells in 30 μL Matrigel per droplet were seeded into a prewarmed 24-well plate. Cells were cultured in DMEM containing 10% FBS, B27 and N2 supplements. The medium was replaced every other day. Sphere size and number were analyzed using ImageJ software.

#### DRG-spheroid coculture

We first built a Matrigel dome containing ACKP cells as described previously. The dome was inverted placed in the incubator for 20 minutes to be solidified. Fresh DRG was placed 1cm or 1 mm from the Matrigel dome with sterile forceps and then covered by 20 μL Matrigel droplet. The plate was inverted placed in the incubator for another 20 minutes. After Matrigel polymerization, the well was filled with RPMI-1640 medium containing GlutaMAX supplement, antibiotic-antimycotic solution, and 10% FBS. To activate sensory neurons, CNO (10 μM) were added in the medium. Images were taken using a Leica Stellaris 8 Confocal Microscope with TokaiHit thermoincubator for live cell imaging.

#### Calcium imaging

In vitro calcium imaging was done with the DRG-spheroid coculture model. We utilized rAAV-CMV-NES-jRGECO1a (10^11^ vg/mL, Biohippo) to infect gastric cancer spheroids. After 7 days infection, spheroids were cocultured with DRG from Trpv1-Cre;hM3Dq;R26-tdTomato mouse. CNO (10 μM) was added to activate sensory neurons and calcium flux in gastric cancer spheroids was detected using a Leica Stellaris 8 Confocal Microscope with TokaiHit thermoincubator.

In vivo calcium imaging was done with the Trpv1-Cre;GCaMP6s;tdTomato mouse. First, ACKP gastric cancer cells were infected with pLV-ChR2 lentivirus (VectorBuilder) to express Channel Rhodopsin. Then, on Day 1, 1 million ACKP-ChR2 cells were injected into the stomach wall of Trpv1-Cre;GCaMP6s;tdTomato mouse to establish gastric orthotopic allograft. On Day 14, the mouse was anesthetized with isoflurane, fixed in a head holder in supine position. We exposed the nodose ganglia by blunt dissection and imaged it on a Leica TCS SP5 upright confocal microscope in XYZT mode. The temporal resolution is 3 second, and spatial resolution is 512 x 512 pixels. Then we opened the abdominal wall. At time 0, we irradiated the gastric tumor by 473 nm blue light. Meanwhile, GCaMP6s fluorescence was continuously recorded at 488Ex/510-550Em.

#### Rabies tracing of cancer-innervating neurons

Monosynaptic rabies neurotracing was performed as previously ^27,55^. Briefly, ACKP gastric cancer cells were infected with LV-EF1a-TVA-P2A-RVG lentivirus (Biohippo) to express TVA and RVG. Then, on Day 1, 1 million ACKP-TVA-RVG cells were injected into the stomach wall of C57BL/6 mouse to establish gastric orthotopic allograft. On Day 14, the mouse was anesthetized with isoflurane, EnvA-ΔG-dsRed rabies virus (10^8^ IFU/mL, 10 μL/mouse, Biohippo) was injected into the gastric cancer. On day 28, mice were sacrificed and nodose ganglia and dorsal root ganglia (T3-L3) were dissected.

#### E2F activity measurement

1 x 10^6^ ACKP cells were transfected with 3 µg pE2F-TA-luc plasmid (Takara) and 0.3 µg renilla luciferase control plasmid pRL-CMV (Promega) using lipofectamine 3000. 24 hours later, ACKP cells were cultured in serum-free medium and treated with CGRP or kinase inhibitors for 24 hours. Cells were harvested and luciferase activity of cell lysates was measured using the Dual-Luciferase Reporter Assay System (Promega) in a GloMax 20/20 Luminometer (Promega) according to manufacturer instructions.

#### Statistics

Statistical analysis was performed with SPSS software (version 25.0). Parametric data were analyzed using two-tailed unpaired Student’s t-tests for two groups. Datasets containing more than two groups were compared using one-way analysis of variance and Dunnett’s test. For non-parametric data, the Mann-Whitney U test was used. Spearman’s correlation test was used to evaluate the correlation between CGRP and p-Rb. The probability of differences in OS was ascertained using the Kaplan-Meier method, with a log-rank test probe for significance. P<0.05 was considered to indicate a statistically significant difference.

### DATA AVAILABILITY

The accession number for the sequencing data reported in this paper is GSE243578.

