## Extended data for "Nociceptive neurons interact directly with gastric cancer cells via a CGRP/Ramp1 axis to promote tumor progression"

**Extended Data Fig. 1**

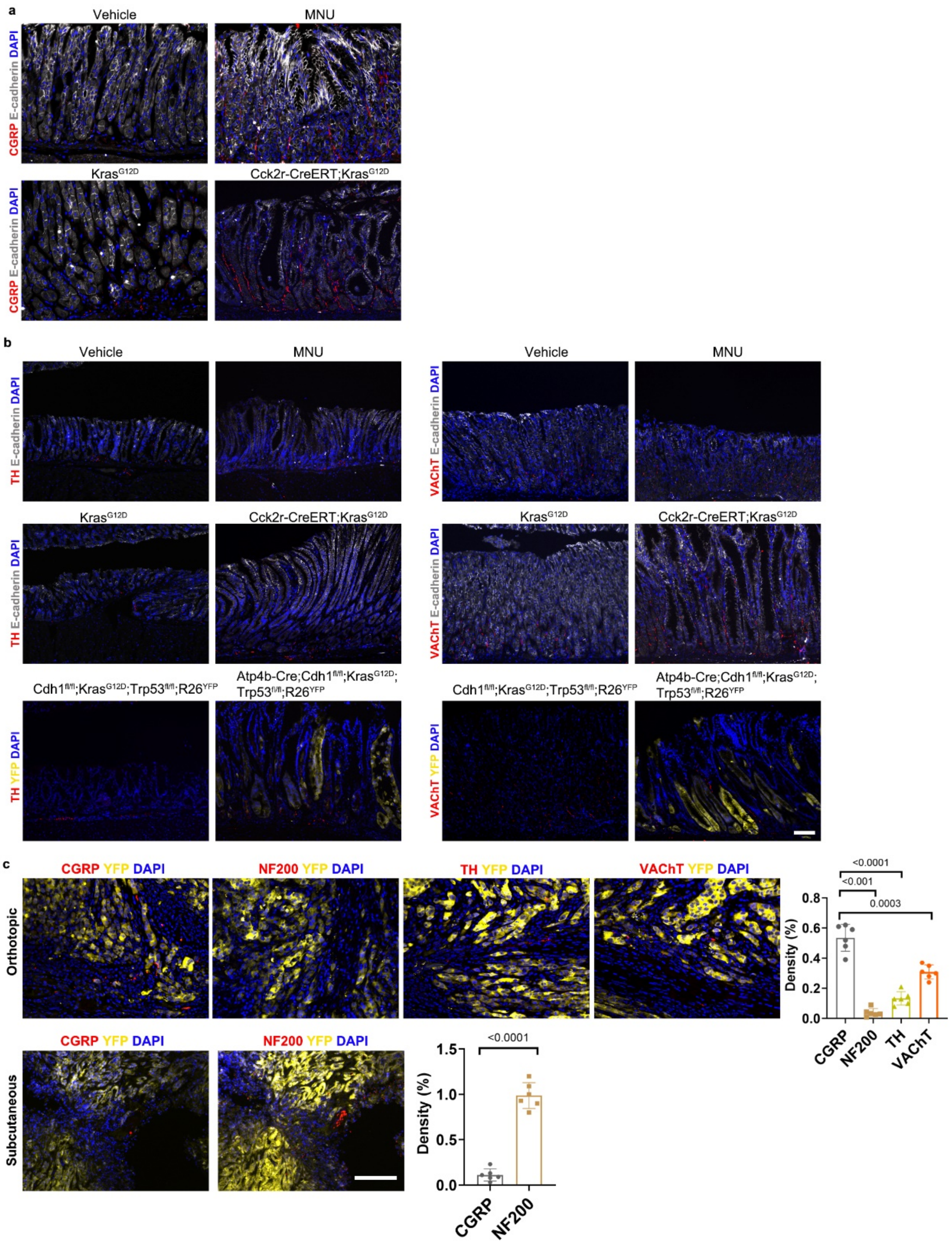


**Neural expansion in GCs.** Representative images of **(a)** sensory (CGRP), **(b)** sympathetic (TH) and parasympathetic (VAChT) nerves in mouse gastric cancers (n = 5/group). Scale bar, 100 μm. (**c**) Representative images and quantification of nociceptive nerves (CGRP), proprioceptive/mechanoreceptive nerves (NF200), sympathetic (TH) and parasympathetic (VAChT) nerves in mouse orthotopic and subcutaneous tumors (n = 6/group). Scale bar, 100 μm. Data represent mean ± SEM, and *P* values were calculated by ANOVA in **c**.

### Extended Data Fig. 2

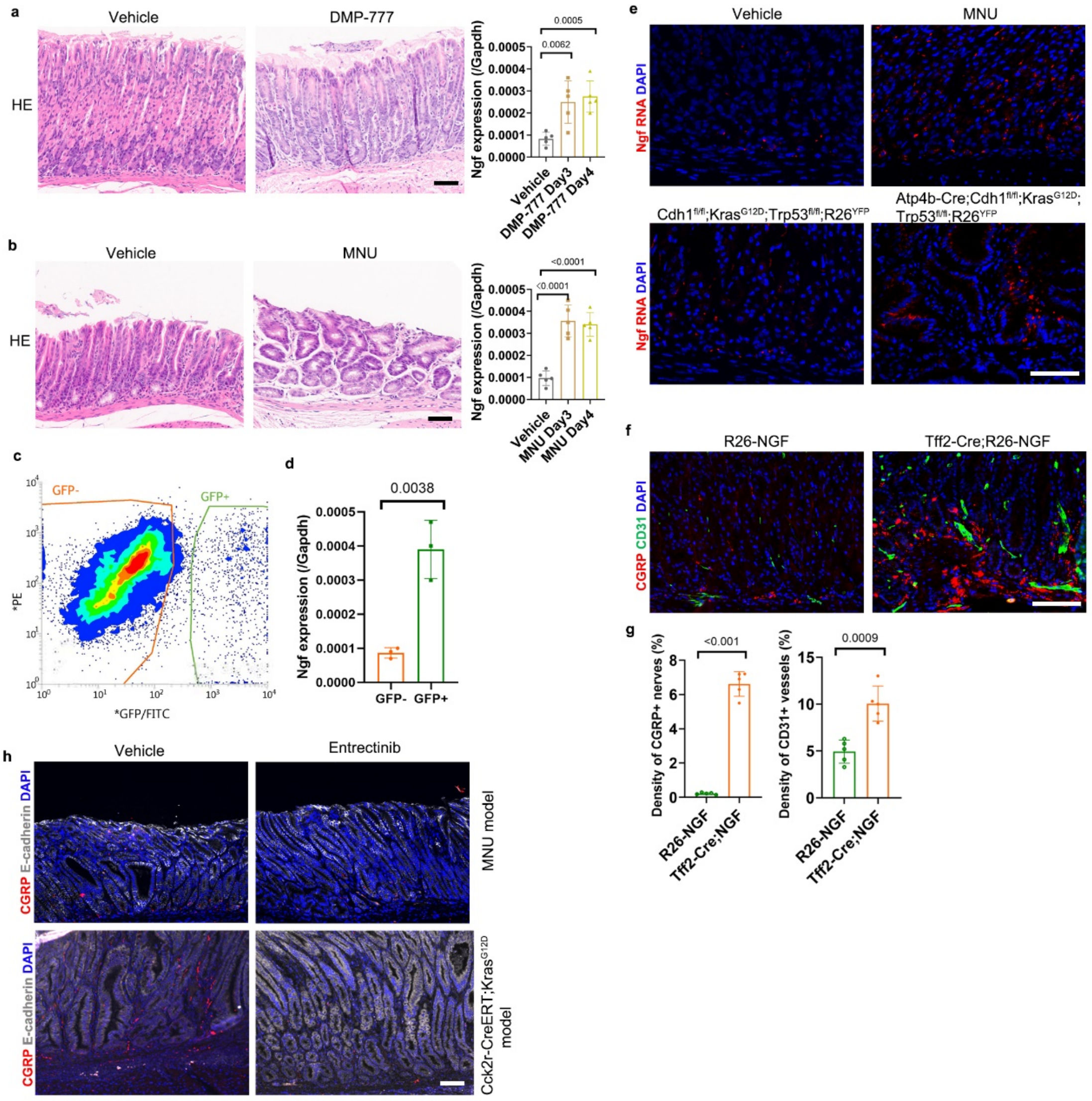


**Gastric epithelial injure upregulated Ngf expression.** (**a**) Representative images of HE staining in mouse stomach treated with vehicle or DMP-777, and Ngf expression in epithelial cells (n = 5/group). Scale bar, 100 µm. (**b**) Representative images of HE staining in mouse stomach treated with vehicle or high dose of MNU, and Ngf expression in epithelial cells (n = 5/group). Scale bar, 100 µm. (**c**) Mist1+ Kras-mutant gastric epithelial cells were isolated from *Mist1*-CreERT; Kras$^{G12D}$; tGFP mice after tamoxifen induction 1 month. (**d**) Relative expression of Ngf in non-mutant and Kras-mutant gastric epithelial cells (n = 3/group). Scale bar, 100 µm. (**e**) Representative images of Ngf in situ hybridization in mouse GCs (n = 5/group). Scale bar, 100 µm. (**f**) Representative images and (**g**) quantification of sensory nerves and micro vessels in NGF-overexpression mice and control mice (n = 5/group). Scale bar, 100 µm. (**h**) Representative images of sensory nerves in mouse gastric cancers treated by Entrectinib or Vehicle (n = 5/group). Scale bar, 100 µm. Data represent mean ± SEM, and *P* values were calculated by ANOVA in **a** and **b**, by t test in **d** and **g**.

**Extended Data Fig. 3**

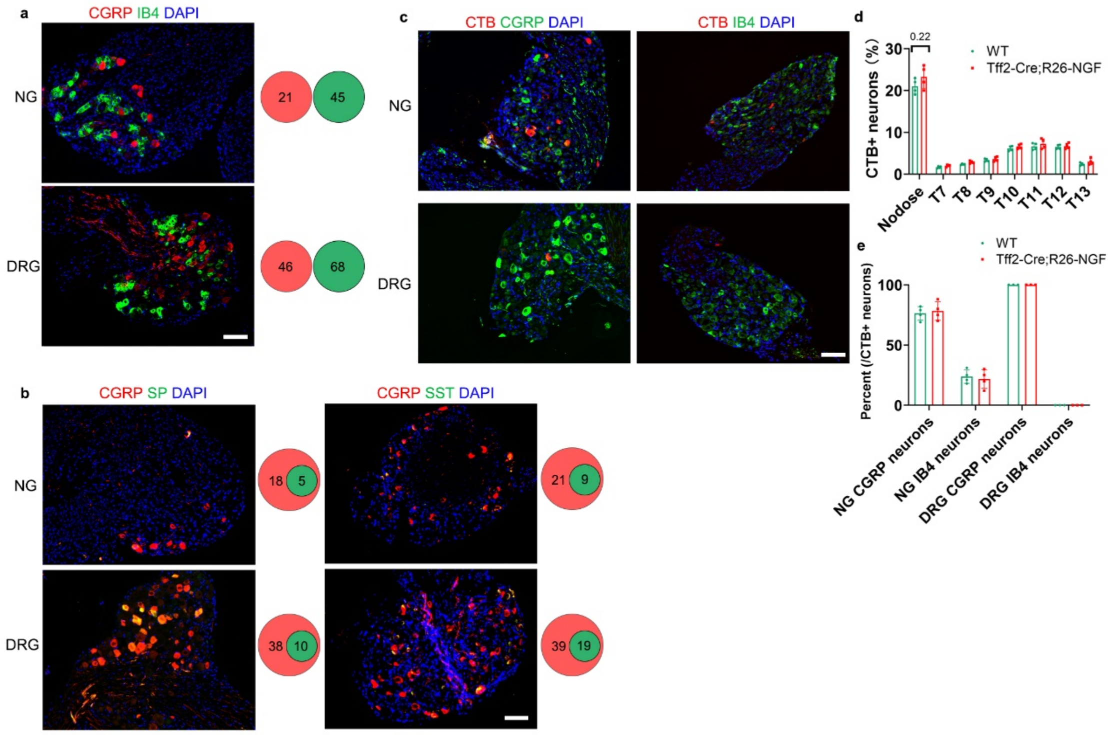


**Nociceptive neurons in ganglia.** (**a**) Representative images and quantification of CGRP+ neurons and IB4+ neurons in ganglia. Scale bar, 100 µm. (**b**) Representative images and quantification of CGRP+, SP+, SST+ neurons in ganglia. Scale bar, 100 µm. SP, Substance P. SST, Somatostatin. (**c**) Representative images and (**d**) quantification of neural tracing in NGF- overexpression mouse (n = 4/group). Scale bar, 100 µm. (**e**) Quantification of molecular subtypes of the stomach-innervating sensory neurons in NGF- overexpression mouse (n = 4/group). Data represent mean ± SEM, and *P* values were calculated by t test.

**Extended Data Fig. 4**

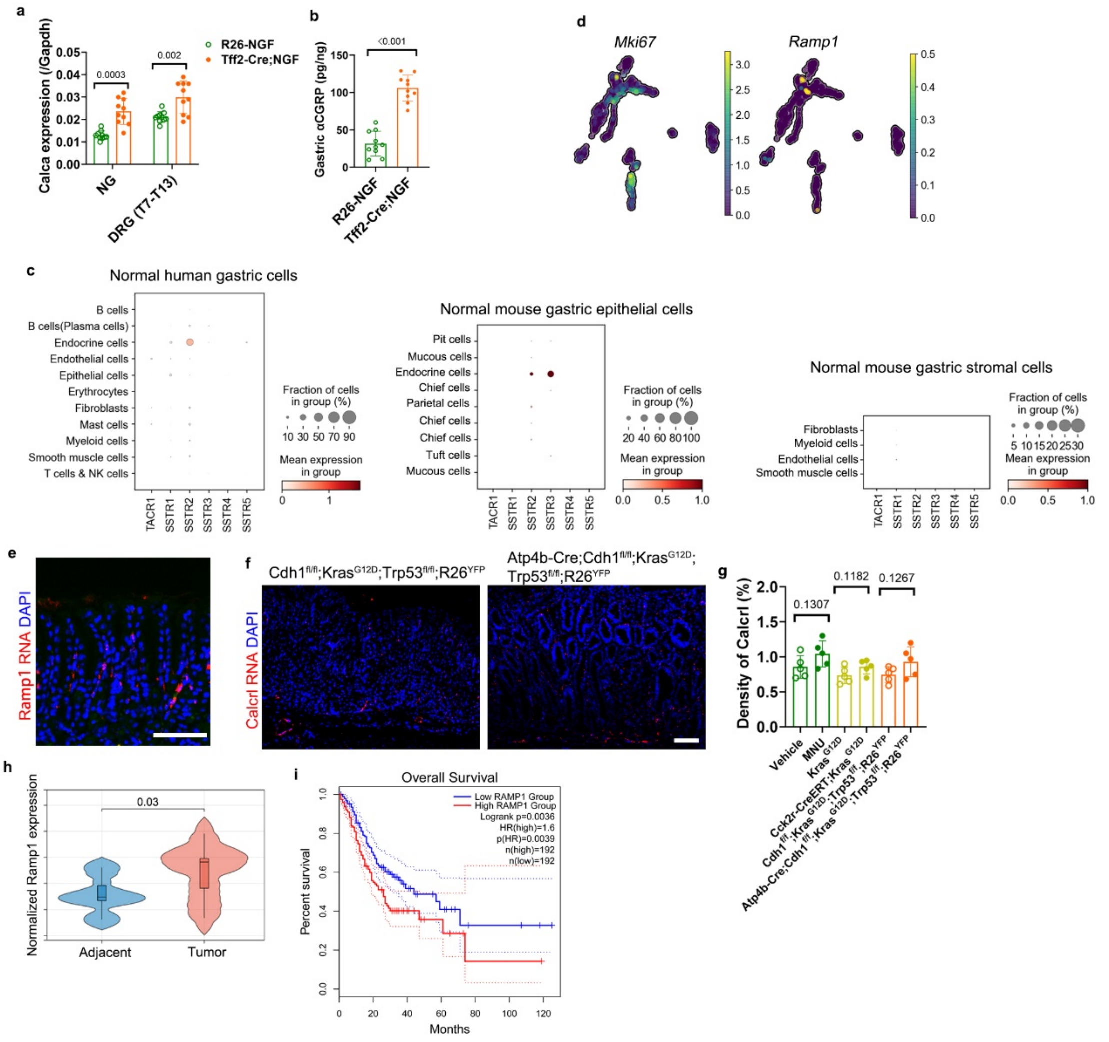


**CGRP/Ramp1 expression in GCs.** (**a**) Expression levels of *Calca* in the stomach-innervating sensory neurons from the NGF-overexpression mice and the control mice (n = 10/group). (**b**) Concentrations of CGRP peptide in the stomach from the NGF-overexpression mice and the control mice (n = 10/group). (**c**) Dot plot of single-cell transcriptome data of Substance P receptors and Somatostatin receptors from mouse stomach (GSE157694 and GSE116514) and human stomach (OMIX001073). (**d**) UMAP of single-cell transcriptome data from mouse stomach (GSE157694). (**e**) Representative image of in situ hybridization of Ramp1 RNA in mouse gastric antrum (n = 5/group). Scale bar, 100 µm. (**f**) Representative images and (**g**) quantification of in situ hybridization (RNAScope) of Calcrl RNA in mouse stomach and gastric cancers (n = 5/group). Scale bar, 100 µm. (**h**) Violin plot of bulk RNA sequencing data from human GC and adjacent tissue (OMIX001073). (**i**) Kaplan-Meier curve of GEPIA survival data. GC patient cohort data were split based on high and low RAMP1 expression. Data represent mean ± SEM, and *P* values were calculated by t test in **a**, **b**, **g** and **h**, by Logrank in **i**.

**Extended Data Fig. 5**

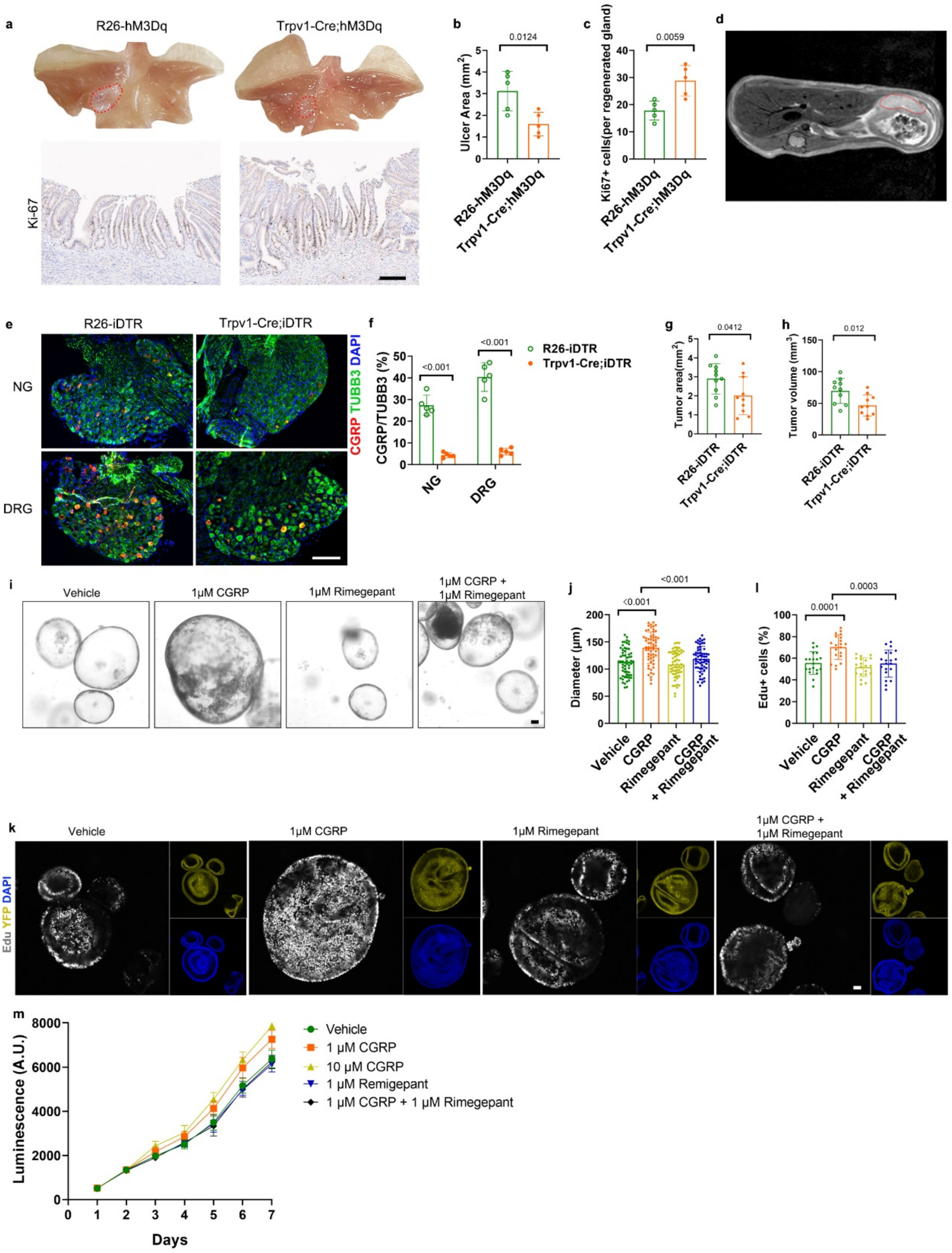


**Nociceptive neurons enhanced gastric ulcer regeneration and cancer cell proliferation.** (**a**) Representative images of gastric ulcer samples and Ki-67 staining (n = 5/group). Scale bar, 100 μm. Quantification of (**b**) ulcer area and (**c**) Ki-67 staining. (**d**) Representative MRI images from orthotopic GC model. (**e**) Representative images and (**f**) quantification of DT-induced nociceptive neuron ablation (n = 5/group). Scale bar, 100 μm. (**g**) Quantification of MNU-induced GCs from nociceptive neuron-ablated mice or control mice (n = 10/group). (**h**) Quantification of syngeneic orthotopic tumors from nociceptive neuron-ablated mice or control mice (n = 10/group). (**i**)

Representative images and (**j**) quantification of gastric cancer spheroids treated by CGRP or Rimegepant. Scale bar, 100 µm. (**k**) Representative images and (**l**) quantification of Edu staining in (**i**). Scale bar, 100 µm. (**m**) Proliferation curve of ACKP cells (n = 3/group). Data represent mean ± SEM, and *P* values were calculated by t test in **b**, **c**, **f**, **g**, **h**, **j** and **l**.

**Extended Data Fig. 6**

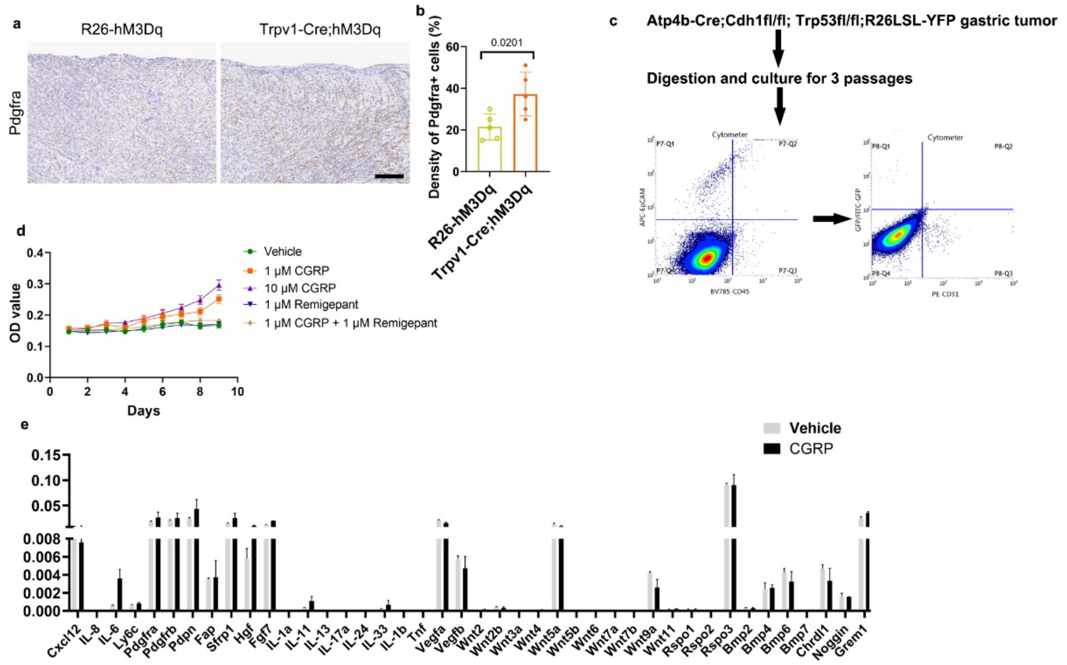


**Nociceptive neurons regulated the number and function of gastric CAFs.** (**a**) Representative images and (**b**) quantification of Pdgfra+ CAFs in orthotopic tumors (n = 5/group). Scale bar, 100 µm. (**c**) Flow chart of gastric CAFs isolation. (**d**) Proliferation curve of gastric CAFs (n = 3/group). (**e**) Expression levels of CAF-associated cytokines in gastric CAFs treated with vehicle of CGRP. Data represent mean ± SEM, and *P* values were calculated by t test in **b**.

**Extended Data Fig. 7**

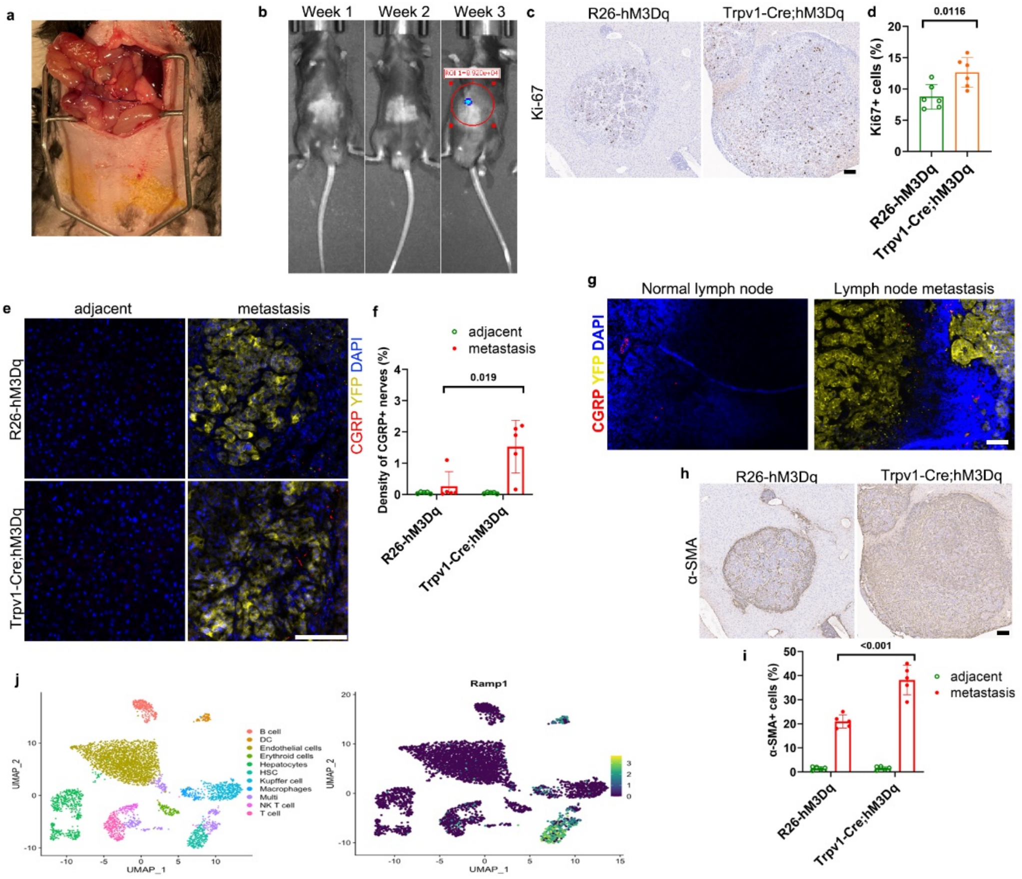


**Nociceptive neurons promote GC metastasis in a spontaneous metastasis model.** (**a**) The spontaneous metastatic model was done by resecting the primary tumor followed by an esophagojejunostomy with Roux-en-Y anastomosis. (**b**) Mice were monitored weekly with bioluminescence (IVIS) following resection of the primary tumor. Liver metastases were detectable from the third week. (**c**) Representative images and (**d**) quantification of Ki-67 staining in liver metastasis. Scale bar, 100 μm. (**e**) Representative images and (**f**) quantification of sensory nerves (red) in liver metastatic area (yellow) and adjacent area. Scale bar, 100 μm. (**g**) Representative images of sensory nerves in normal and metastatic lymph nodes. Scale bar, 100 μm. (**h**) Representative images and (**i**) quantification of α-SMA staining in liver metastasis. Scale bar, 100 μm. (**j**) UMAP of single-cell transcriptome data from mouse liver (GSE174748). Data represent mean ± SEM, and *P* values were calculated by t test in **d, f** and **i**.

**Extended Data Fig. 8**

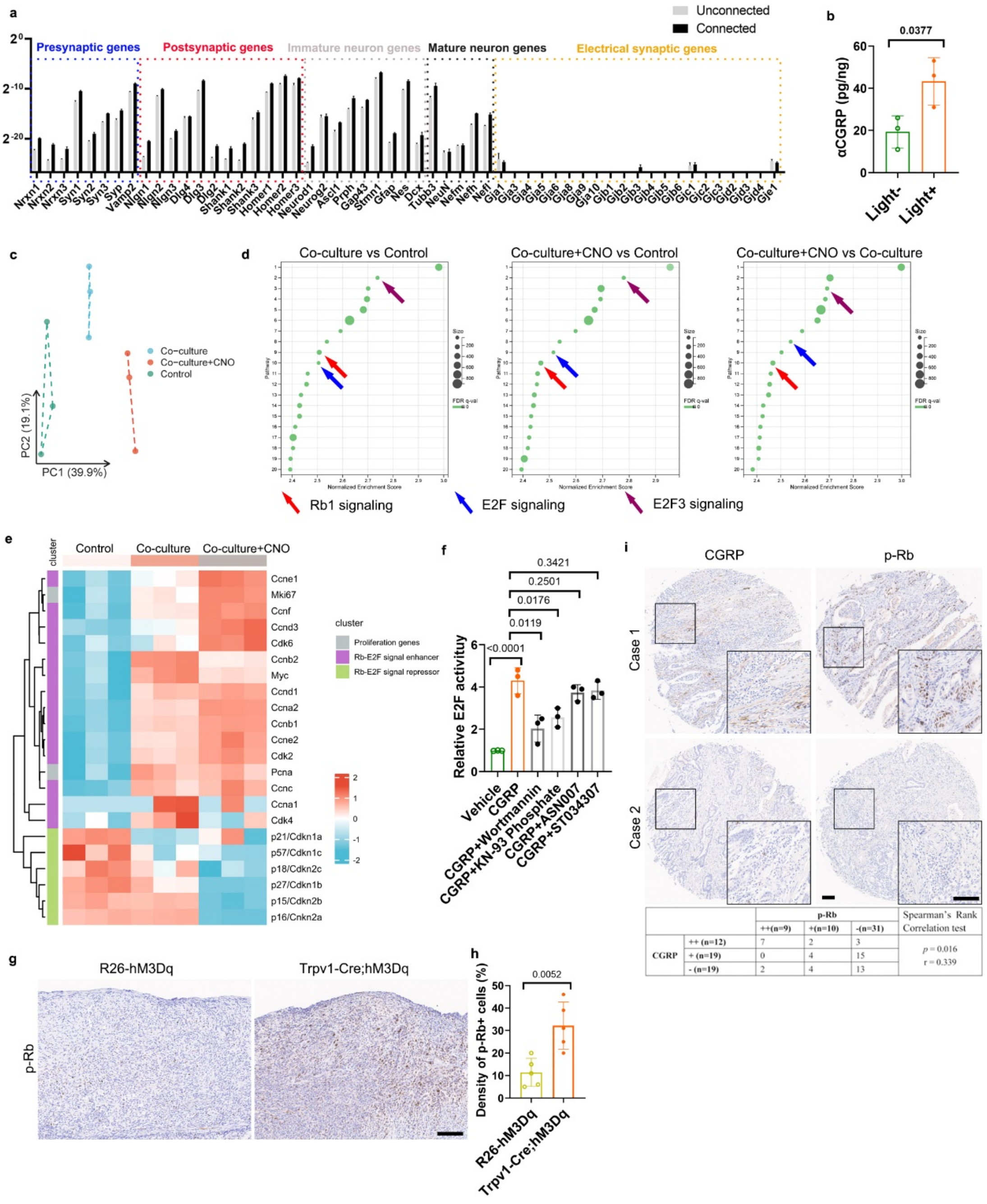


**Nociceptive neurons activate Rb/E2F signaling depending on CaMK and PI3K.** (**a**) Neuron-connected spheroids and unconnected spheroids were split. Expression levels of chemical synaptic genes, electrical synaptic genes and neurodevelopmental genes were analyzed. (**b**) Concentrations of CGRP peptide in orthotopic ACKP-ChR2 tumors from *Trpv1*-Cre; GCamp6s; tdTomato mice (n = 3/group). (**c**) PCA plot of bulk RNA sequencing data from cancer spheroids alone, cancer spheroids cocultured with DRG, and cancer spheroids cocultured with CNO-activated DRG (n = 3/group). (**d**) GSEA plot of top 20 enhanced pathways in each comparison. (**e**) Heat map of proliferation genes, Rb-E2F signal enhancer genes and Rb-E2F signal repressor genes. (**f**) Relative E2F activity in ACKP cells treated with CGRP or kinase inhibitors. Wortmannin, PI3K inhibitor. KN-93, CaMK inhibitor. ASN007, ERK inhibitor. ST034307, PKA inhibitor. (**g**) Representative images and (**h**) quantification of p-Rb staining in orthotopic tumors from CNO-treated mice (n = 5/group). Scale bar, 100 μm. (**i**) Representative images and correlation test of sensory nerve and p-Rb staining in human GC tissue microarray (n = 50). Scale bar, 100 μm. Data represent mean ± SEM, and *P* values were calculated by ANOVA in **f**, by t test in **b** and **h**, and by spearman's rank correlation test in **i**.

**Extended Data Fig. 9**

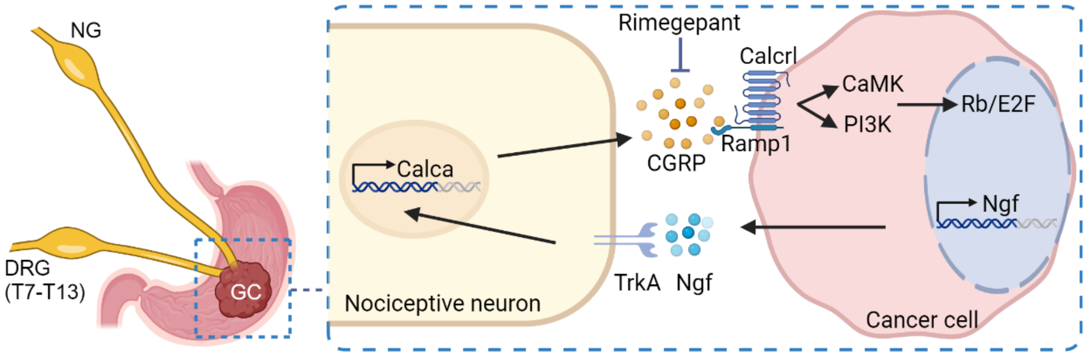


**Graphical abstract.** Gastric cancer is innervated by nociceptive neurons from ipsilateral nodose ganglia and dorsal root ganglia (T7-T13). Gastric cancer cells increase the expression of NGF which attracts the expansion of nociceptive nerves. NGF binds to TrkA receptor causing CGRP synthesis and release from nociceptive nerves. In turn, CGRP activates Calcrl/Ramp1 receptor and promotes E2F activity through PI3K signaling and CaMK signaling in gastric cancer cells.
